# Seroprevalence of zoonotic diseases (leptospirosis and Japanese encephalitis) in swine in ten provinces of Vietnam: a Bayesian approach to estimate prevalence

**DOI:** 10.1101/584151

**Authors:** Hu Suk Lee, To Long Thanh, Nguyen Khanh Ly, Hung Nguyen-Viet, Krishna K. Thakur, Delia Grace

## Abstract

**Background:** Leptospirosis is an important zoonotic disease with a global distribution, affecting a wide range of mammalian animals and humans. Japanese encephalitis (JE) virus is the major vector-borne zoonotic disease in the Asia-Pacific region. The main objective of this study was to evaluate the seroprevalence of serovar-specific *Leptospira* and JE in swine from 10 provinces in Vietnam.

**Methods:** Samples were initially collected for swine influenza surveillance from March to April 2017 at large-scale farms (with at least 50 sows and/or 250 fattening pigs) with pigs that tested positive for influenza in the previous surveillance period (2015-16).

**Findings:** A total of 2,000 sera samples were analyzed from 10 provinces. Overall, the seroprevalence of leptospirosis was 21.05% (95% CI: 19.28-22.90) using a cut-off titer of ≥ 1:100. The apparent prevalence of JE was 73.45% (95% CI: 71.46-75.37) while the true prevalence was slightly higher (74.46%, 95% credible interval: 73.73-86.41). We found a relatively high presence of leptospirosis and JE in pigs kept on large farms. Prevalence was comparable with other studies suggesting opportunistic testing of samples collected for other surveillance purposes can be a valuable tool to better understand and prevent the potential transmission of these zoonotic diseases from pigs to people in Vietnam.

**Conclusion:** Our study provides evidence to veterinarians and animal health professionals for evidence-based practice such as diagnosis, vaccination and zoonotic control. Further investigation into the possible role of different domestic animals, wildlife species or environmental factors is needed to identify the potential risk factors and transmission routes in Vietnam.

## 1. Introduction

Leptospirosis is an important zoonotic disease with a global distribution, affecting a wide range of mammalian animals and humans. Japanese encephalitis (JE) virus is the major vector-borne zoonotic disease in the Asia-Pacific region[1–5]. *Leptospira* spp. are maintained in a wide range of reservoir hosts, such as cattle, pigs, dogs, rats, raccoons, skunks and opossums[6–10]. Leptospirosis is caused by gram-negative bacteria, and more than 260 pathogenic serovars have been identified[11]. Pigs are considered to be reservoir hosts for serovars Bratislava, Pomona and Tarassovi Mitis, and also become infected with Icterohaemorrhagiae from rats and Hardjo from cattle [12–14]. JE virus is a flavivirus and the main cause of viral encephalitis in Asian countries, resulting in 13,600-20,400 deaths per year [15,16]. It is estimated that more than 3 billion people live in endemic areas [17]. JE virus circulates between *Culex* sp. mosquitoes and domestic/wild birds or pigs [18]. JE in pigs can cause abortion, stillbirth and infertility which may result in significant economic losses to producers [19,20]. Pigs are considered to be amplifying hosts and an important risk factor for transmission of JE to humans [21,22]. Bird migration may play an important role in the spread of JEV[15].

In Vietnam, leptospirosis is a notifiable disease in humans, but few cases (less than 20 cases per year) have been reported to the Ministry of Health. Despite the low numbers of reported cases, leptospirosis is considered to be endemic and serovar Bratislava is commonly observed [23–26]. A study in the Mekong delta reported a seroprevalence of 18.8% among 1,400 randomly selected people aged between 15 and 60 years[27]. A study in pigs, using titer ≥1:100 by the microscopic agglutination test (MAT), found a seroprevalence of 73% among 424 sows in the Mekong delta area[23]. The serovar Bratislava showed the highest followed by Autumnalis.

JE virus was first isolated in 1951 in Vietnam, and then it was detected in birds, pigs and in the late 1960s and 1970s[28,29]. A study to evaluate JE virus infection in mosquitoes in Vietnam from 2006 to 2008 showed that several JE virus genotype I populations were circulating across the country[30]. A recent study found a seroprevalence was 60.4% in 641 samples of pig sera[31]. A JE vaccine was introduced in 1997 and is used for children under five years of age as part of Vietnam’s national immunization program in 12 high-risk diseases in the northern region[4].

To our knowledge, only a few multi-site studies have been implemented to assess the seroprevalence of leptospirosis and JE in swine and these have covered in limited areas in Vietnam. Therefore, the main objective of this study was to evaluate the seroprevalence of serovar-specific *Leptospira* and JE in swine from 10 provinces in Vietnam.

## 2. Material and methods

### 2.1 Study locations and sampling

Samples were initially collected for swine influenza surveillance from March to April 2017 under the Department of Animal Health (DAH) and Food and Agriculture organization (FAO) of the United Nations. The Department selected 10 provinces (Ha Noi, Bac Ninh, Thai Binh, Bac Giang, Quang Ninh, Quang Ngai, Binh Duong, Dong Nai, Dong Thap and Soc Trang) (Fig. 1). The sampling was targeted at large-scale farms (with at least 50 sows and/or 250 fattening pigs) with pigs that tested positive for influenza in the previous surveillance period (2015-16). In each province, 10 blood samples from piglets (4-8 weeks old) and fattening pigs (9-12 weeks old) and five blood samples from sows were collected from four-large scale farms, but the sex of the pigs was not recorded. Pigs with symptoms of influenza were prioritized for sampling.

**Figure 1.**
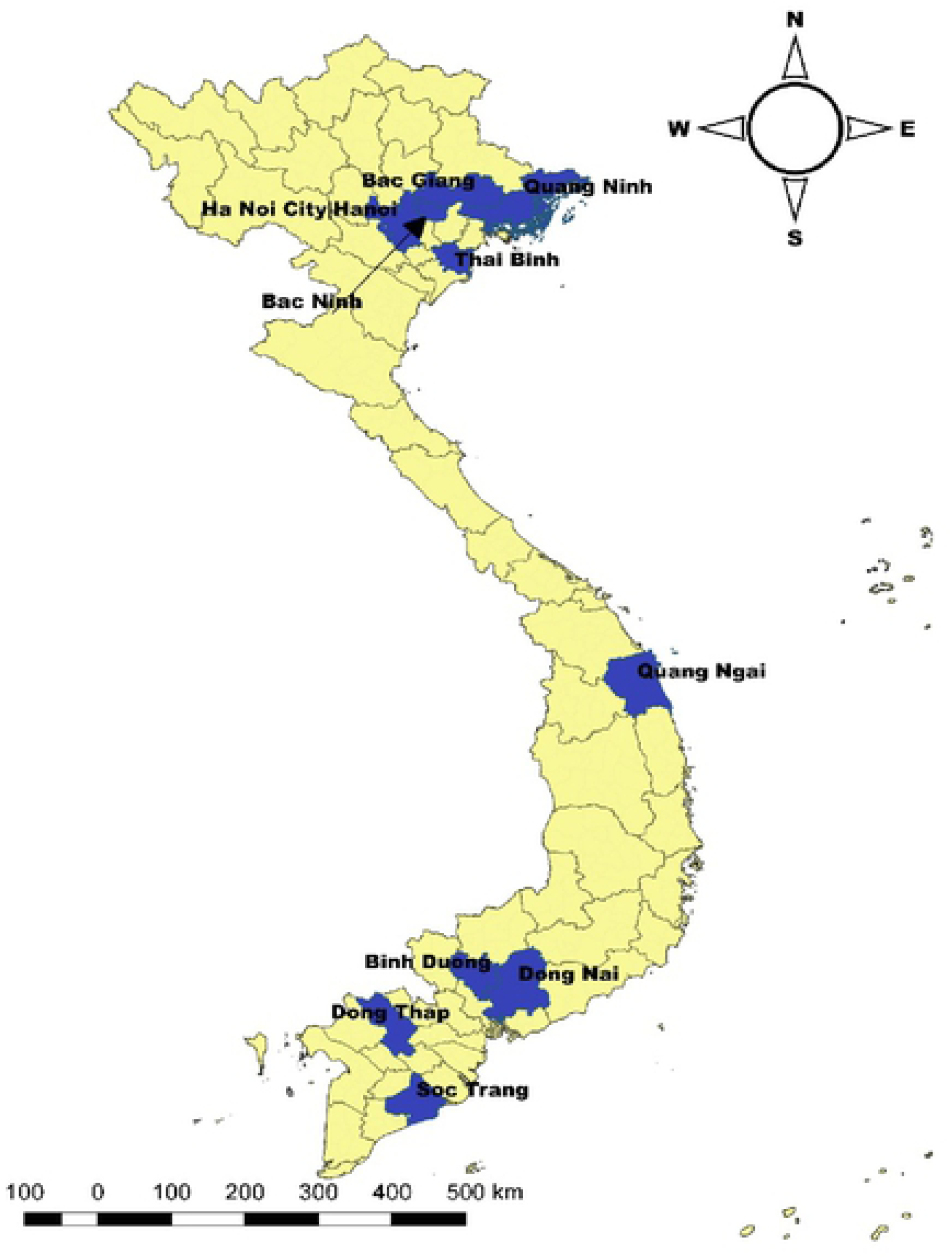
Selected sampling provinces (in blue) for leptospirosis and JE in swine.

### 2.2 Laboratory analysis

The sera were extracted after centrifugation and stored at −20 °C until transportation in a cool box to the National Center for Veterinary Diagnosis (NCVD) in Hanoi. For leptospirosis, the MAT was used to assess the positivity of samples. Two-fold serial dilution was used after an initial 1:100 dilution, and the end-point (1:1600) was the highest dilution of serum that shows at least 50% agglutination of live leptospires compared to the control sample. For JE, enzyme-linked immunosorbent assay (ELISA) was used to detect JE virus-specific antibodies in swine serum. We followed the manufacturer’s instructions of JE Ab ELISA protocol (VDPro® JE Ab ELISA; Median, Chuncheonsi, Korea). A sample was determined as positive if the S/P ratio 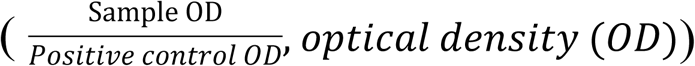 is more than 0.25.

### Data analysis

We considered a sample leptospirosis positive if the MAT was ≥ 1:100 for at least one of the 16 serovars (table 1). For JE, we estimated the true prevalence (TP) from apparent prevalence (AP) by adjusting for the diagnostic test sensitivity and specificity using a Bayesian framework [32]. The prior distribution for sensitivity [uniform(0.953-0.987)] and specificity [uniform (0.95-0.953)] were obtained from the company protocol, and a non-informative Jeffreys prior [beta = (0.5, 0.5)] was used to minimize the influence on the posterior [33]. Markov Chain Monte Carlo (MCMC) sampling in “truePrev” was conducted by JAGS through the rjags package [34,35]. The first 1,000 samples of the three MCMC chains were discarded as a burn-in period and the following 10,000 iterations were used for posterior inference. The AP was calculated based on the proportion of positive samples with a 95% confidence interval (CI) while the TP was estimated based on the posterior median value with a 95% credible interval. The outputs from the three chains were assessed visually using MCMC trace-plots, posterior density distribution plots, Brooks-Gelman-Rubin (BGR) plots, and auto-correlation plots using the CODA package. All data were entered in Microsoft Excel 2016 and analyzed using R (version 3.5.2). QGIS (Quantum GIS development Team 2018. QGIS version number 3.0.1) was used to generate the map.

**Table 1.** List of *Leptospira* antigens used in the MAT.

| No. | Genomospecies | Serogroup | Serovar |
| --- | --- | --- | --- |
| 1 | *L. interrogans* | Australis | Australis |
| 2 | *L. interrogans* | Autumnalis | Autumnaliss |
| 3 | *L. interrogans* | Bataviae | Bataviae |
| 4 | *L. interrogans* | Australis | Bratislava |
| 5 | *L. interrogans* | Canicola | Canicola |
| 6 | *L. kirschneri* | Grippotyphosa | Grippotyphosa |
| 7 | *L. interrogans* | Hebdomadis | Hebdomadis |
| 8 | *L. interrogans* | Icterohaemorrhagiae | Icterohaemorrhagiae |
| 9 | *L. borgpetersenii* | Javanica | Javanica |
| 10 | *L. noguchii* | Panama | Panama |
| 11 | *L. interrogans* | Pomona | Pomona |
| 12 | *L. interrogans* | Pyrogenes | Pyrogenes |
| 13 | *L. borgpetersenii* | Sejroe | Hardjo |
| 14 | *L. borgpetersenii* | Sejroe | Saxkoebing |
| 15 | *L. biflexa* | Semarang | Patoc |
| 16 | *L. borgpetersenii* | Tarassovi | Tarassovi |

Source: http://leptospira.amc.nl/leptospira-library/leptospira-strains/?grid-page=2

## 3. Results

### 3.1 Leptospirosis

A total of 2,000 sera samples were analyzed from 10 provinces. Overall, the seroprevalence of leptospirosis was 21.05% (95% CI: 19.28-22.90) using a cut-off titer of ≥ 1:100 (Table 2). Quang Ngai showed the highest seroprevalence (37.5%) followed by Binh Duong (32.5%) whereas Soc Trang had the lowest sero-prevalence (10.0%), and the differences were statistically significant. By using a low cut-off titer (≥ 1:100), the most frequently observed presumptive infective serovar was Bratislava (9.65%), followed by Pyrogenes (6.1%) and Tarassovi (4.55%) (table 3). Using a cut-off titer (≥ 1:200), Bratislava (1.95%) and Panama (0.75%) had the highest seroprevalences. By province, Bratislava had the highest sero-prevalence of the 8 provinces (except Hanoi and Quang Ngai) (Fig. 2). Serovar Tarassovi and Pyrogenes had the highest seroprevalences in Hanoi and Quang Ngai, respectively. A total of 144 samples were positive with more than two serovars, which was accounting for 34.20% (95% CI: 29.68-38.95) among positive samples. Two samples from Binh Dung and Dong Nai were positive with maximum seven serovars. The sero-prevalence of sow (42.78%, 95% CI: 37.85-47.83%) was significantly higher than younger age groups (5-8 weeks and 9-12 weeks) (Fig. 3).

**Table 2.** Sero-prevalence (%) with 95% CI for *Leptospira* serovars in pigs in Vietnam using MAT.

| Province (no.) | Sero-positive samples<br>(a titer≥1:100 for any serovars) | Sero-positive (%) with 95% CI |
| --- | --- | --- |
| Bac Giang (200) | 61 | 30.5 (24.20-37.39) |
| Bac Ninh (200) | 45 | 22.5 (16.91-28.92) |
| Binh Duong (200) | 65 | 32.5 (26.06-39.47) |
| Dong Nai (200) | 38 | 19.0 (13.81-25.13) |
| Dong Thap (200) | 23 | 11.5 (7.43-16.75) |
| Hanoi (200) | 23 | 11.5 (7.43-16.75) |
| Quang Ngai (200) | 75 | 37.5 (30.77-44.61) |
| Quang Ninh (200) | 43 | 21.5 (16.02-27.85) |
| Soc Trang (200) | 20 | 10.0 (6.22-15.02) |
| Thai Binh (200) | 28 | 14.0 (9.51-19.59) |
| Total (2,000) | 421 | 21.05 (19.28-22.90) |

**Table 3.** MAT results for Leptospira serovars in pigs by using 3 cutoff titers

| Serovar | Total samples | ≥ 1:100 | ≥ 1:200 | ≥ 400 |
| --- | --- | --- | --- | --- |
|  |  | N (% , 95% CI) | N (% , 95% CI) | N (% , 95% CI) |
| Australis | 2,000 | 45 (2.25, 1.65-3.00) | 4 (0.2, 0.005-0.05) | 0 |
| Autumnalis | 2,000 | 1 (0.05, 0.0001-0.28) | 0 | 0 |
| Bataviae | 2,000 | 10 (0.5, 0.24-0.92) | 1 (0.05, 0.0001-0.28) | 0 |
| Bratislava | 2,000 | 193 (9.65, 8.39-11.03) | 39 (1.95, 1.39-2.66) | 8 (0.4, 0.17-0.79) |
| Canicola | 2,000 | 45 (2.25, 1.65-3.00) | 2 (0.1, 0.01-0.36) | 0 |
| Grippotyphosa | 2,000 | 18 (0.9, 0.53-1.42) | 2 (0.1, 0.01-0.36) | 0 |
| Icterohaemorrhagiae | 2,000 | 22 (1.1, 0.69-1.66) | 3 (0.15, 0.03-0.44) | 0 |
| Javanica | 2,000 | 13 (0.65, 0.35-1.11) | 4 (0.2, 0.005-0.05) | 0 |
| Panama | 2,000 | 87 (4.35, 3.50-5.34) | 15 (0.75, 0.42-1.23) | 2 (0.1, 0.01-0.36) |
| Pomona | 2,000 | 31 (1.55, 1.06-2.19) | 3 | 0 |
| Pyrogenes | 2,000 | 122 (6.1, 5.09-7.24) | 8 (0.4, 0.17-0.79) | 0 |
| Hardjo | 2,000 | 0 | 0 | 0 |
| Sakoebing | 2,000 | 0 | 0 | 0 |
| Tarassovi | 2,000 | 91 (4.55, 3.68-5.56) | 8 (0.4, 0.17-0.79) | 1 (0.05, 0.0001-0.28) |
| Patoc | 2,000 | 5 (0.25, 0.08-0.58) | 1 (0.05, 0.0001-0.28) | 0 |

**Figure 2.**
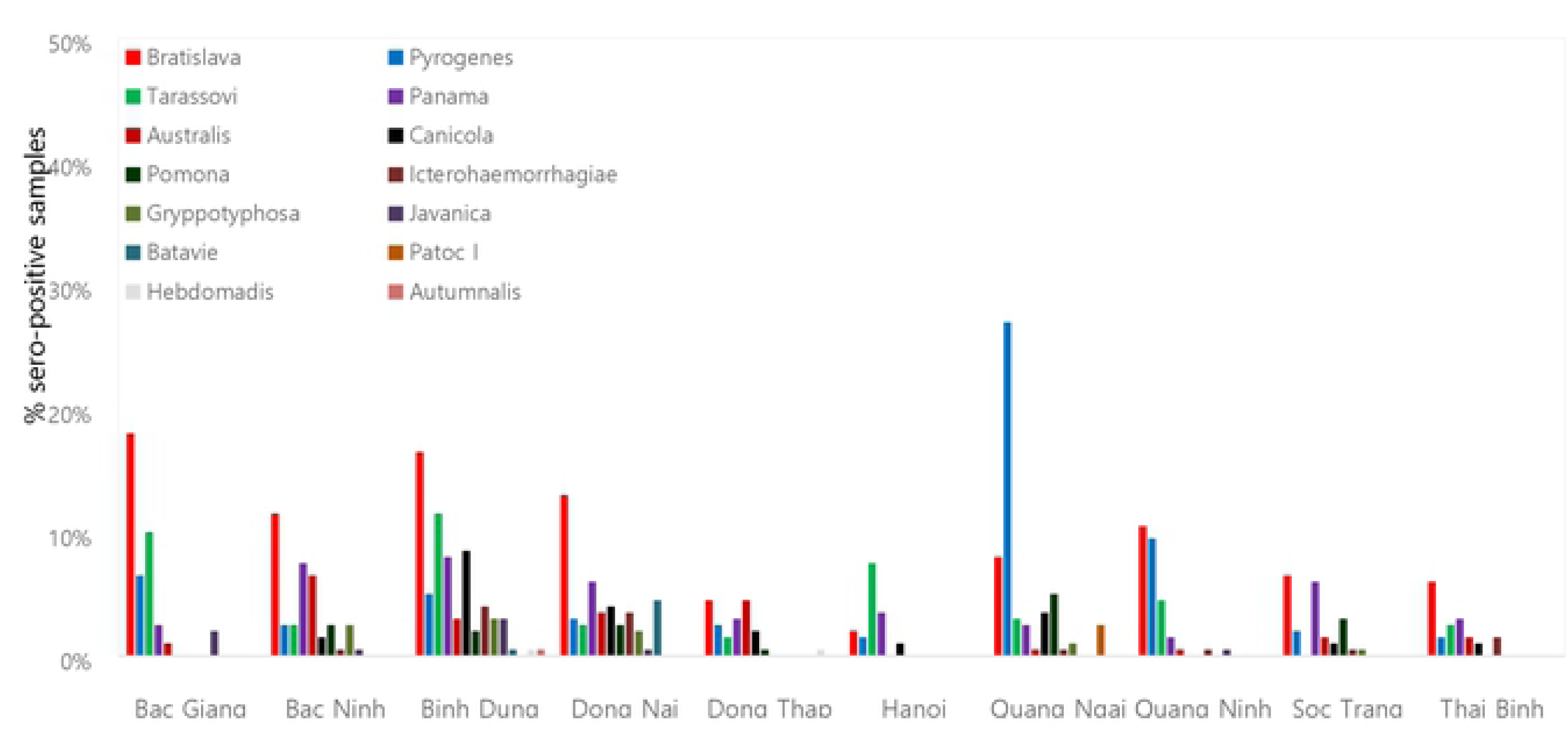
Production of sero-positive samples by serovar in each province using cutoff titer ≥ 1:100.

**Figure 3.**
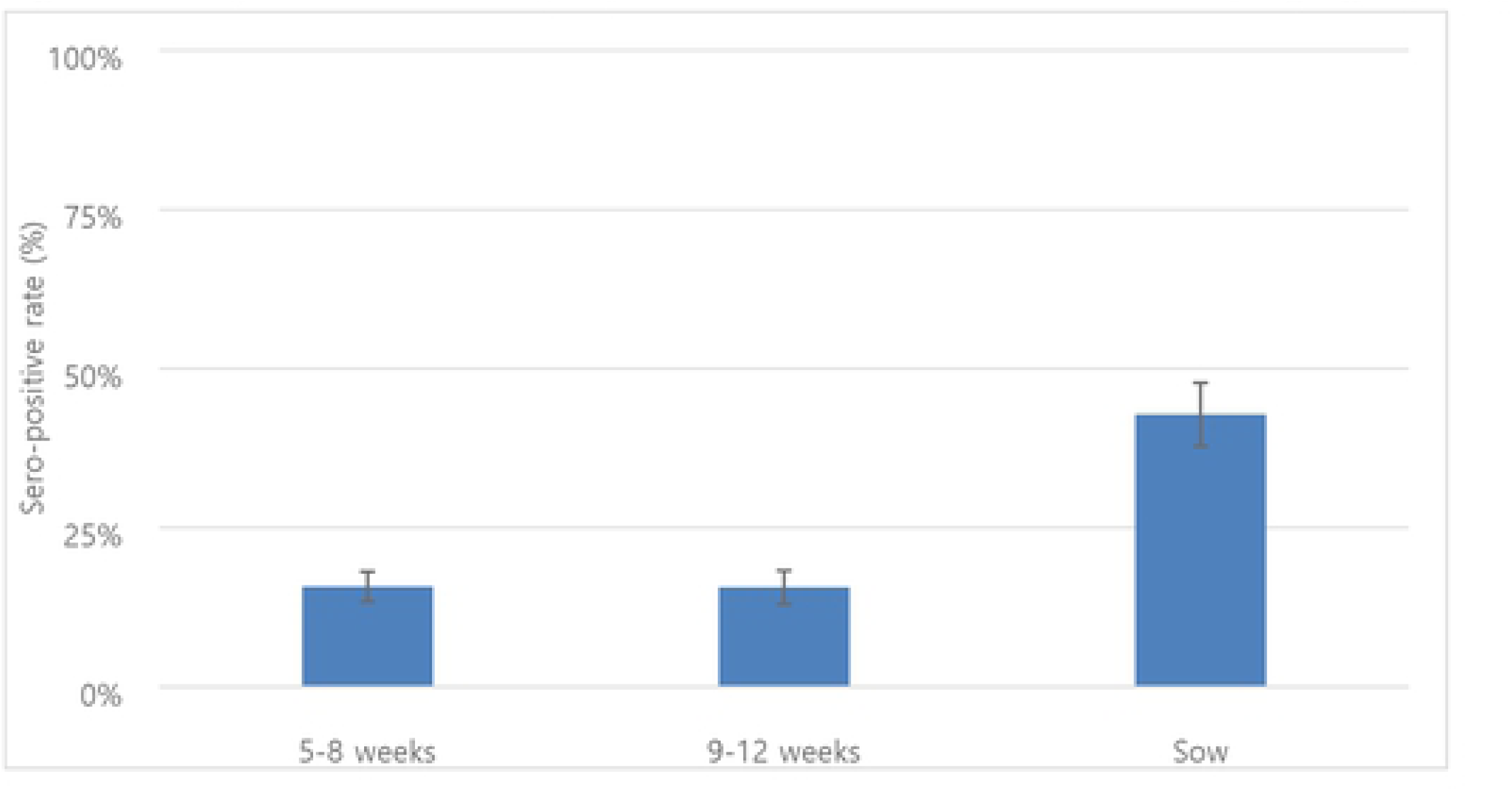
Seropositive rates of leptospirosis with 95% confidence interval by age group in pigs using cut off titer ≥ 1:100.

### 3.2 Japanese encephalitis (JE)

Overall, the AP of JE was 73.45% (95% CI: 71.46-75.37) while the TP was slightly higher (74.46%, 95% credible interval: 73.73-86.41) (Table 4). Dong Thap had the highest seroprevalence followed by Thai Binh whereas Binh Dung showed the lowest seroprevalence. Overall, the AP showed slightly higher than the TP except for Binh Duong province. The seroprevalence in sows (82.27%, 95% CI: 78.15-85.92) was significantly higher than that in animals of other age groups (Fig. 4). The BGR plots showed that our chains have converged for TP, sensitivity and specificity, respectively (Fig. 5).

**Table 4.** Apparent sero-prevalence with 95% CI and true sero-prevalence with 95% credible interval for Japanese encephalitis in pigs in Vietnam.

| Province (no.) | Sero-positive samples | Apparent sero-prevalence (%)<br>with 95% CI | True sero-prevalence (%)<sup>a</sup><br>with 95% credible interval |
| --- | --- | --- | --- |
| Bac Giang (200) | 158 | 79.0 (72.69-84.43) | 80.47 (73.73-86.41) |
| Bac Ninh (200) | 141 | 70.5 (63.66-76.72) | 71.21 (63.97-77.96) |
| Binh Duong (200) | 122 | 61.0 (53.87-67.80) | 60.92 (53.20-68.25) |
| Dong Nai (200) | 155 | 77.5 (71.08-83.09) | 78.83 (71.96-84.88) |
| Dong Thap (200) | 168 | 84.0 (78.17-88.79) | 85.93 (79.73-91.36) |
| Hanoi (200) | 125 | 62.5 (55.39-69.23) | 62.55 (55.14-69.70) |
| Quang Ngai (200) | 128 | 64.0 (56.93-70.65) | 64.22 (56.66-71.37) |
| Quang Ninh (200) | 161 | 80.5 (74.32-85.75) | 82.12 (75.58-87.89) |
| Soc Trang (200) | 145 | 72.5 (65.76-78.56) | 73.46 (66.30-79.94) |
| Thai Binh (200) | 166 | 83.0 (77.06-87.93) | 84.79 (78.46-90.39) |
| Total (2,000) | 1,469 | 73.45 (71.46-75.37) | 74.46 (71.84-77.06) |

<sup>a</sup> Median value was recorded

**Figure 4.**
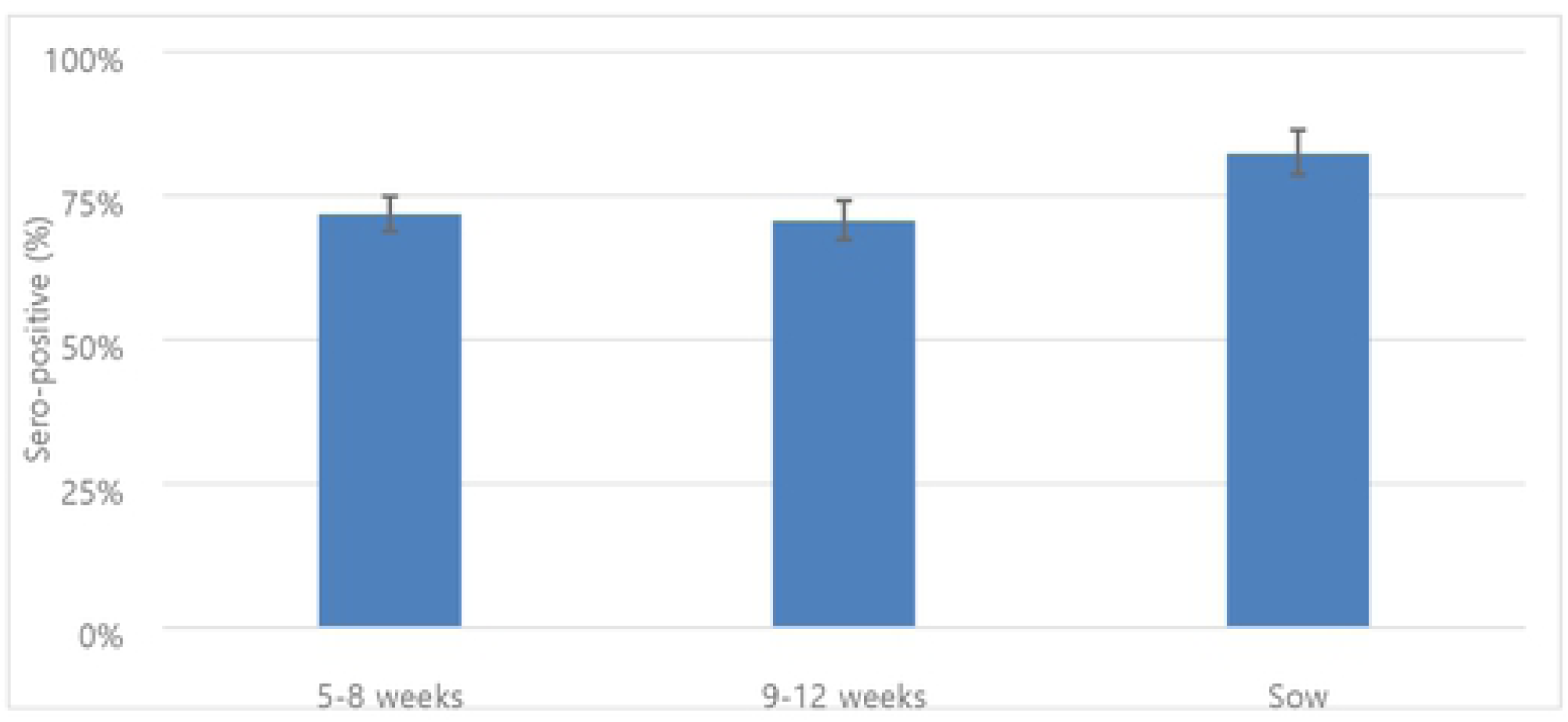
Seropositive rates of japanese encephalitis with 95% confidence interval by age group in pigs.

**Figure 5.**
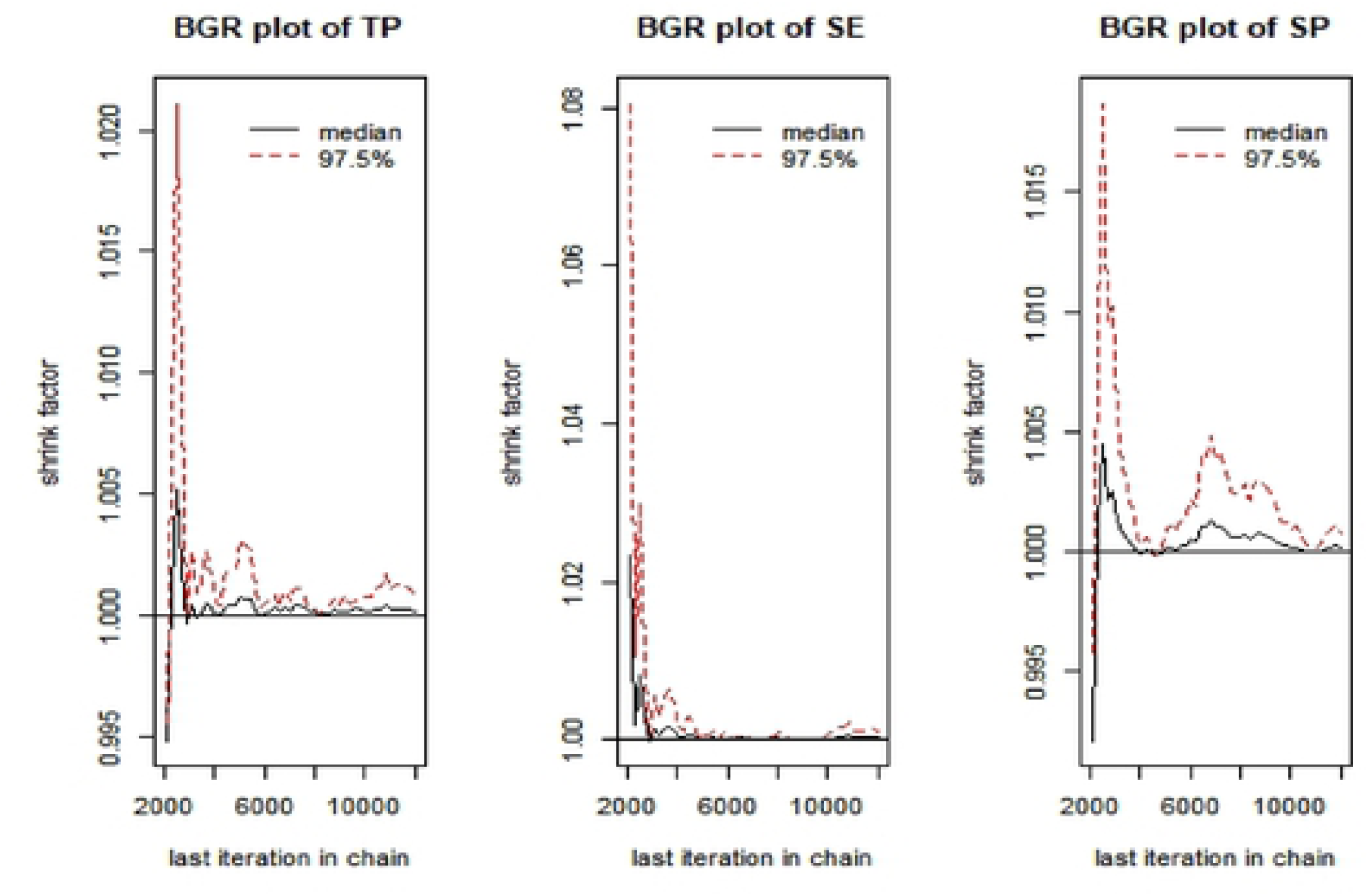
Brooks-Gelman-Rubin (BGR)plots for True prevalence (TP), Sensitivity (SE) and Specify.

## 4. Discussion

A nation-wide study on seroprevalences of leptospirosis and JE in pigs was conducted in Vietnam. Overall, the seroprevalence (21.05%; 95% CI: 19.28-22.90) of leptospirosis was higher than that reported by Lee et al. (8.17%)[36]. A possible explanation is that pig samples from that study were randomly collected from slaughterhouses and it may be that healthy or less visually ill pigs were more likely to be sent for slaughter. Other studies conducted in the early 2000s reported higher seroprevalences (26%-73%) using a cut-off titer of ≥ 1:100 [23,37]. These study sites were mostly in the Mekong Delta, close to river areas, so that pigs in these areas may have more opportunities exposure. However, in our study, samples from three provinces located in the Mekong Delta had a lower seroprevalence [(Soc Trang (10.0%), Dong Thap (11.5%) and Dong Nai (19.0%)] than other provinces. Therefore, further investigation is necessary to assess the association between environmental factors and leptospirosis.

Our study found that Bratislava and Tarassovi Mitis were more common, similar to the previous studies [23,36]. Mostly, pigs are considered to be the main host for serovars Tarassovi Mitis, Pomona, Bratislava and Muenchen [13,38,39]. Other studies found that Bratislava, Pomona and Tarassovi Mitis were commonly detected in wild boars [40,41]. Therefore, there is a possibility of leptospirosis transmission from wild boars to domesticated pigs. Further study is needed to better understand the role of wild animals in the transmission of leptospirosis in Vietnam. Serovars Pyogenes and Panama were newly detected in our study, which have been observed in cattle in other countries [42–44]. Interestingly, significantly higher prevalence of serovar Pyrogenes was detected in Quang Ngai. According to the Vietnam General statistics office, Quang Ngai has the third largest cattle population in the country (approximately 277,000), after Nghe An and Gia Lai (all central areas) [45], where was only included from the central region for our study. Therefore, it would be worthwhile to conduct an epidemiological investigation on the circulation of serovar Pyrogenes in the central region, where the largest cattle population is raised in Vietnam. Moreover, Icterohaemorrhagiae and Javanica were observed that rat may play an important role in the transmission to pigs [46]. One study suggests that older pigs have a higher chance of exposure to *Letospira* which is consistent with our result [23]. In humans, leptospirosis is one of 28 notifiable diseases in Vietnam. A total of 85 cases were officially recorded between 2008 and 2015. The low number of notifications is due to lack of public awareness and diagnostic facilities in rural areas resulting in under-reporting. Therefore, it is important to raise public awareness, especially among high-risk groups (e.g. agriculture/livestock farmers, mining workers, veterinarians and slaughterhouse workers) who are more likely to come in contact with infected animals and contaminated water or soil.

Overall, the seroprevalence (73.45%) of JE in pigs was similar to that in neighboring countries (Laos and Cambodia [65-75%]) [47,48], but higher than that in some other Asian countries (Indonesia, Nepal and Taiwan) [49–52]. In Vietnam, pigs were not vaccinated against JE virus mainly due to economic reasons, implying that positivity was due to natural infection.

Furthermore, seroprevalence (82%) in pigs older than two months showed that detected antibodies were caused by virus infections rather than maternal immunity (one study found that maternal antibodies disappeared after two months [53]). Therefore, we can conclude that JE is widespread in the pig population across the country. In Southeast Asia, it is well documented that JE is endemic and a public health concern. Current pig production practices may pose a risk for JE, because people live very close to pigs, increasing exposure to mosquitoes with JE virus infection. In Vietnam, small-scale backyard pig production accounts for 70-75% of the total production[54].

The average seroprevalence from five northern provinces was 75.1% (95% CI: 72.42-77.78) which was significantly higher than the previous study (60.4% among 641 pig sera), sampling conducted in the northern part of Vietnam from 2009 to 2010 [31]. However, there is no evidence that JE is increasing over time in Vietnam. Over the last decade, in Vietnam, livestock production is undergoing a major change, a moving toward intensive systems and large production scales [55]. Therefore, a possible explain is the growth of intensive pig farms which provides more opportunities to encounter mosquitoes with JE virus infection.

In humans, most cases are in unvaccinated children under14 years of age[56]. JE vaccine was first introduced in 1997 through the national immunization program for children aged 1–5 years and 85% of districts were covered in 2013. Currently, there are no routine pig national surveillance programs for JE virus. Therefore, it is difficult to assess the potential role of pigs in the transmission cycle to humans as well as the impact of the national immunization program. We found less discrepancy between them as sensitivity and specificity are relatively high (> 95%) as well as a higher proportion of positive sample. A Bayesian approach provides an opportunity to combine prior information with investigated data and estimate values for both prevalence and diagnostic characteristics of tests[57]. If a diagnostic test with less than 100% sensitivity and specificity is used to estimate the prevalence of a disease, our results would be biased. Therefore, a Bayesian approach was used to estimate the TP from AP. In general, Bayesian models are sensitive to the selection of the priors, so we employed a Jeffreys prior to minimize the influence of prior to the posterior distribution as no prior information was available for JE prevalence in our study area. Our study was the first attempt to estimate the TP for JE in Vietnam, which showed how a Bayesian approach can be used to better estimate the prevalence of animal diseases.

This study shows the added value of opportunistic use of samples collected for one purpose in providing valuable information on other diseases. An advantage is the large sample size and wide cover. A disadvantage is potential bias linked with sample collection from large farms with previous influenza detection and preference given to apparently sick pigs.

## 5. Conclusions

We found a relatively high presence of leptospirosis and JE in pigs kept on large farms. Prevalence was comparable with other studies suggesting opportunistic testing of samples collected for other surveillance purposes can be a valuable tool to better understand and prevent the potential transmission of these zoonotic diseases from pigs to people in Vietnam. Our study provides evidence to veterinarians and animal health professionals for evidence-based practice such as diagnosis, vaccination and zoonotic control. Further investigation into the possible role of different domestic animals, wildlife species or environmental factors is needed to identify the potential risk factors and transmission routes in Vietnam.

## Abbreviations

JE: Japanese encephalitis;
CI: confidence interval;
TP: true prevalence;
AP: apparent prevalence;
ELISA: enzyme-linked immunosorbent assay;
MCMC: Markov Chain Monte Carlo;
BGR: Brooks-Gelman-Rubin

## Ethics approval and consent to participate

The study was approved by the Hanoi Medical University Institutional Review Board (HMU IRB: no. 00003121), Vietnam.

## Acknowledgments

We thank the Department of Animal Health under the Vietnam Ministry of Agriculture and Rural Development and the Food and Agriculture Organization of the United Nations office in Vietnam for sharing samples. We also thank Tezira Lore of the International Livestock Research Institute for editing the manuscript.

## Authors’ contributions

Conceived and designed the experiments: HSL Performed the experiments: HSL, TLT and NKL Analyzed the data: HSL Wrote the paper: HSL, TLT, NKL, HNV, KKT, and DG. All authors read and approved the final manuscript.

## Availability of data and materials

All datasets supporting our findings are available from the corresponding author on reasonable request

## Consent for publication

Not applicable.

